# Neural Signatures of Conscious Experience During Sleep: A Serial Awakening Study Using High-Density EEG

**DOI:** 10.64898/2026.08.16.745040

**Authors:** Atakan Selte, Steven E. Haworth, Thomas Vanasse, Tariq Alauddin, Klevest Gjini, Santiago Philibert-Rosas, Cameron J. Brace, Brinda Sevak, Brady Riedner, Mariel Kalkach Aparicio, Giulio Tononi, Melanie Boly, Aaron F. Struck

## Abstract

Identifying neural signatures of consciousness remains a central challenge in neuroscience. Sleep offers a tractable model for comparing brain activity in the presence or absence of subjective experience while minimizing behavioral responsiveness confounds. Using overnight 256-electrode high-density EEG in 140 participants and a serial-awakening paradigm, we analyzed 699 non-rapid eye movement (NREM) sleep stage 2 and 3 awakenings (351 dreaming experience, 348 no experience). Features from the 60s preceding awakening included regional spectral power, lagged-coherence connectivity, graph-theoretic metrics and gamma-to-alpha power ratios. Dreaming experiences were associated with shifts in posterior spectral balance, particularly reduced alpha and delta power and increased gamma-related measures, together with altered large-scale network organization. In participant-level cross-validated machine-learning analyses, all classifiers performed above chance, with the best ensemble model reaching an ROC-AUC of 0.80 and average precision of 0.80. These findings identify reproducible posterior electrophysiological and network-level signatures of conscious states during NREM sleep.

## Introduction

The investigation of consciousness during sleep has long been a prominent focus in neuroscience, offering unique insights into the brain’s varying states of awareness. Sleep provides an ideal framework for examining consciousness, as it encompasses transitions between different levels of awareness across distinct stages.(1–3) Dreaming studies also enable to isolate the neural signatures of consciousness from those of behavioral responsiveness. (2) While previous studies have reported neural signatures of dreaming experiences during sleep—particularly decreased posterior cortical delta power in small cohorts of subjects—these results have not been confirmed in larger cohorts. Furthermore, machine learning approaches applied to electrophysiological signals allow for a broader parameter search to identify optimal features distinguishing conscious from unconscious states. In this study, we aimed to identify high-density EEG features distinguishing between individuals who reported dreaming experiences and those who did no subjective experience after serial awakenings from non-rapid eye movement (NREM) sleep. Participants were awakened from different stages of sleep via an automated auditory stimulus and asked to report on the epoch immediately preceding awakenings. This method, established as both efficient and reliable in probing consciousness during sleep, (4,5) allowed for the collection of multiple subjective reports per individual.

Prior work on dreaming suggested that local spectral dynamics are important.(6–8), especially reductions in low-frequency activity in the posterior cortical regions.(2,9–11). Therefore as part of our machine learning feature selection, we computed band power across delta, theta, alpha, beta, and gamma bands using Welch’s method within anatomically defined posterior, frontal, and control regions.(12,13) While previous studies solely focused on EEG spectral power, connectivity measures may also sensitively probe large-scale network organization features associated with consciousness.(6–9) In this study, we employed lagged coherence to analyze HD-EEG connectivity. From these connectivity matrices, global graph-theoretic metrics including modularity, global efficiency, and rich-club organization were computed to characterize network segregation and integration. Beyond standalone spectral and connectivity features, we also incorporated engineered composite features designed to capture cross-frequency balance. These include gamma-to-alpha power ratios derived from spectral measures. Such features serve as indirect markers of cortical activation relevant to contemporary theories of consciousness. (18–21) By integrating regional spectral features, network topology features, and engineered features within a unified framework, we aimed to develop a first-of-its-kind model that can forecast dreaming experiences during NREM sleep at the level of single awakenings.

## Results

A total of 699 awakenings met inclusion criteria for EEG signal quality and were analyzed (351 CE and 348 NCE), corresponding to a mean of 4.99 awakenings per participant. One CE subject was excluded prior to analysis due to anomalous spectral power >6 standard deviations above the subject with the second highest power, consistent with likely artifact or non-physiological signal contamination. Statistical screening identified several EEG features that differentiated CE from NCE awakenings. The strongest effects were observed in posterior spectral features: posterior alpha power was lower during CE than NCE (r = −0.82; large effect), posterior delta power was also lower during CE (r = −0.78; moderate effect), and posterior gamma power was lower during CE (r = −0.68; moderate effect), all surviving false discovery rate (FDR) correction (Table 1). Engineered cross-frequency features also showed robust separation, with both the posterior gamma/alpha ratio (r = 0.75; FDR q = 0.0045) and frontal gamma/alpha ratio (r = 0.63; FDR q = 0.0201) higher during CE. Regional channel group definitions used for feature extraction are shown in Figure 1.

**Figure 1.**
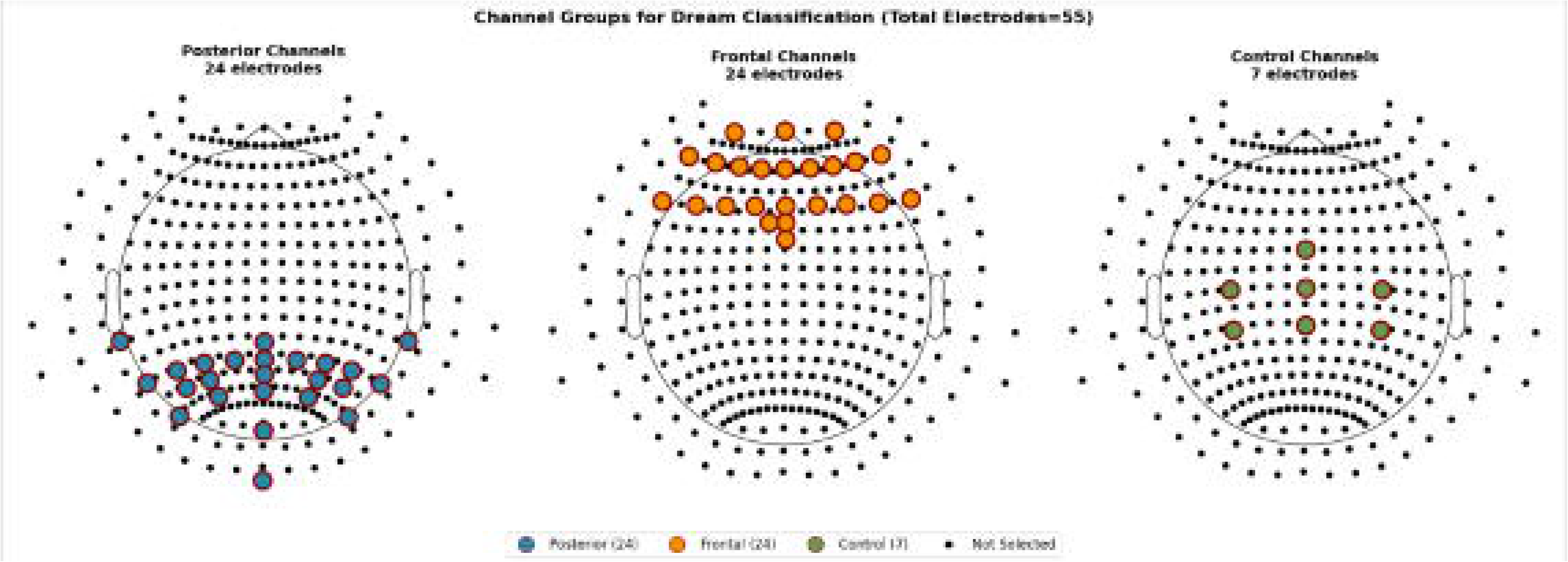
Regional electrode group definitions for feature extraction. High-density EEG electrodes were grouped into posterior (parieto-occipital), frontal (anterior/prefrontal), and control (central) regions to compute regional spectral and engineered cross-frequency features.

**Table 1.**
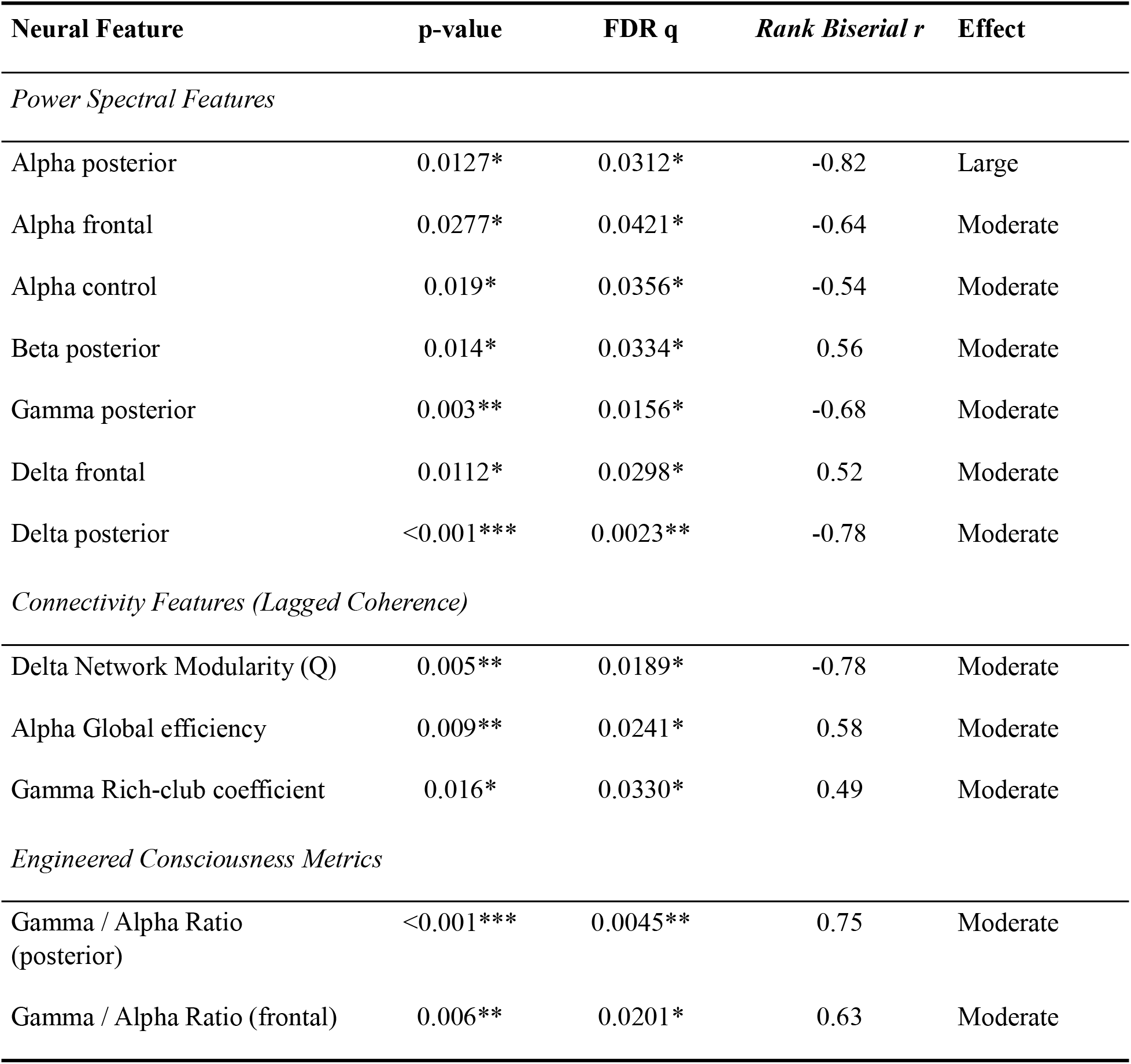
Model Performance Comparison.

| Neural Feature | p-value | FDR q | Rank Biserial r | Effect |
| --- | --- | --- | --- | --- |
| *Power Spectral Features* |  |  |  |  |
| Alpha posterior | 0.0127* | 0.0312* | -0.82 | Large |
| Alpha frontal | 0.0277* | 0.0421* | -0.64 | Moderate |
| Alpha control | 0.019* | 0.0356* | -0.54 | Moderate |
| Beta posterior | 0.014* | 0.0334* | 0.56 | Moderate |
| Gamma posterior | 0.003** | 0.0156* | -0.68 | Moderate |
| Delta frontal | 0.0112* | 0.0298* | 0.52 | Moderate |
| Delta posterior | <0.001*** | 0.0023** | -0.78 | Moderate |
| *Connectivity Features (Lagged Coherence)* |  |  |  |  |
| Delta Network Modularity (Q) | 0.005** | 0.0189* | -0.78 | Moderate |
| Alpha Global efficiency | 0.009** | 0.0241* | 0.58 | Moderate |
| Gamma Rich-club coefficient | 0.016* | 0.0330* | 0.49 | Moderate |
| *Engineered Consciousness Metrics* |  |  |  |  |
| Gamma / Alpha Ratio (posterior) | <0.001*** | 0.0045** | 0.75 | Moderate |
| Gamma / Alpha Ratio (frontal) | 0.006** | 0.0201* | 0.63 | Moderate |

Connectivity-derived features also differentiated CE from NCE awakenings (Table 1, Figure 2). Delta-band network modularity (Q) was lower during CE (r = −0.78), while alpha-band global efficiency (r = 0.58) and gamma-band rich-club coefficient (r = 0.49) were higher during CE, each showing statistically significant group differences after FDR correction. Significant discriminative features spanned spectral, connectivity, and engineered cross-frequency domains

**Figure 2.**
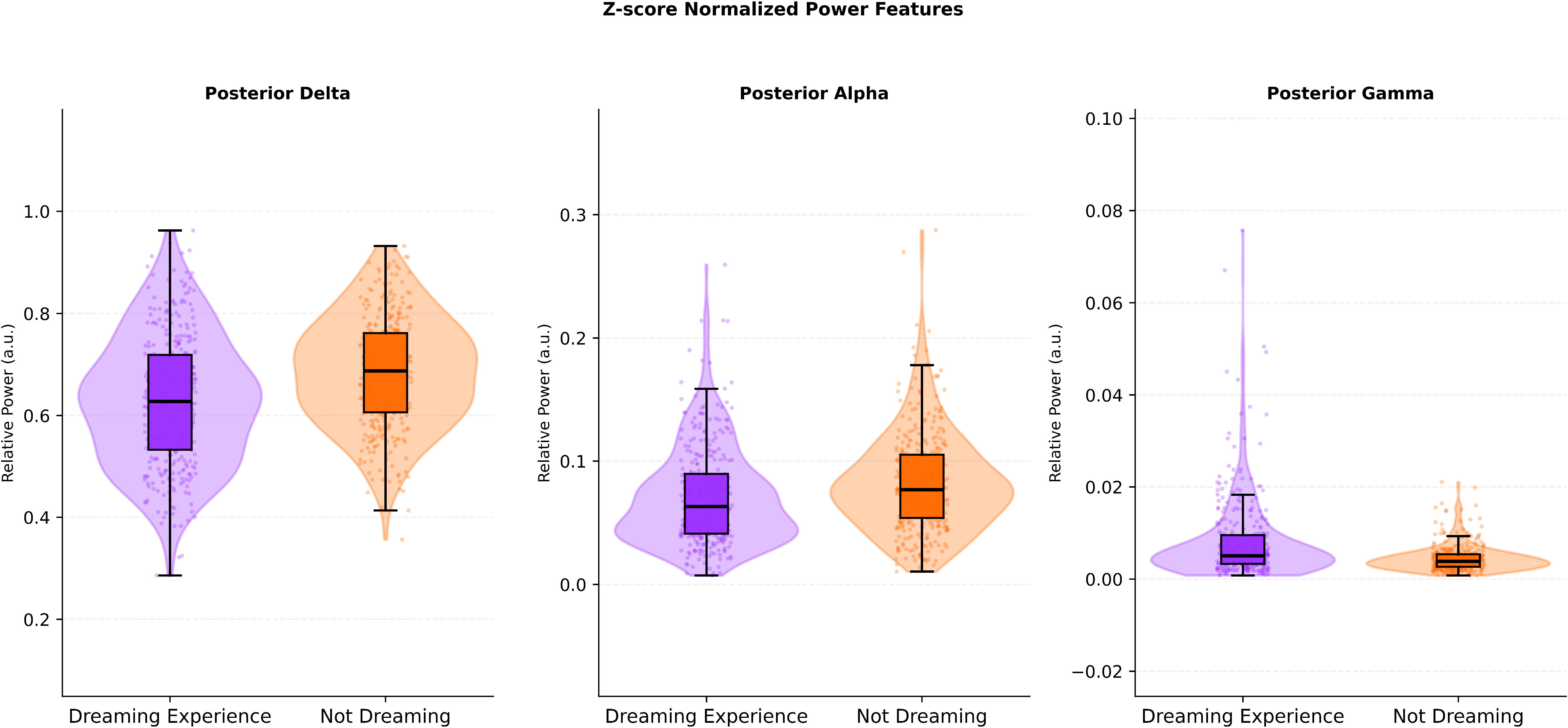
Z-score normalized posterior spectral power during dreaming (conscious experience, CE) versus non-dreaming (no conscious experience, NCE) awakenings. Combined violin and box plots show the distribution of relative power in three posterior frequency bands: delta (left), alpha (middle), and gamma (right). Each point represents a single awakening; box plots indicate the median and interquartile range, with whiskers extending to 1.5× the IQR. Compared with NCE, CE awakenings showed lower posterior delta and alpha power and lower posterior gamma power.

### Classification performance

Selected features were evaluated for their ability to classify CE versus NCE awakenings across multiple supervised learning algorithms (Figure 3). All evaluated classifiers performed above random chance. Discrimination performance measured by ROC-AUC ranged from 0.69 (perceptron) to 0.80 (probabilistic ensemble). Intermediate performance was observed for random forest (AUC 0.76), logistic regression (AUC 0.75), support vector machine (SVM) (AUC 0.74), and k-nearest neighbors (KNN) (AUC 0.71) (Figure 3A). Precision recall performance showed a similar ordering, with the probabilistic ensemble achieving the highest average precision (AP 0.80), followed by random forest (AP 0.74), logistic regression (AP 0.74), SVM (AP 0.72), KNN (AP 0.68), and perceptron (AP 0.65) (Figure 3B).

**Figure 3.**
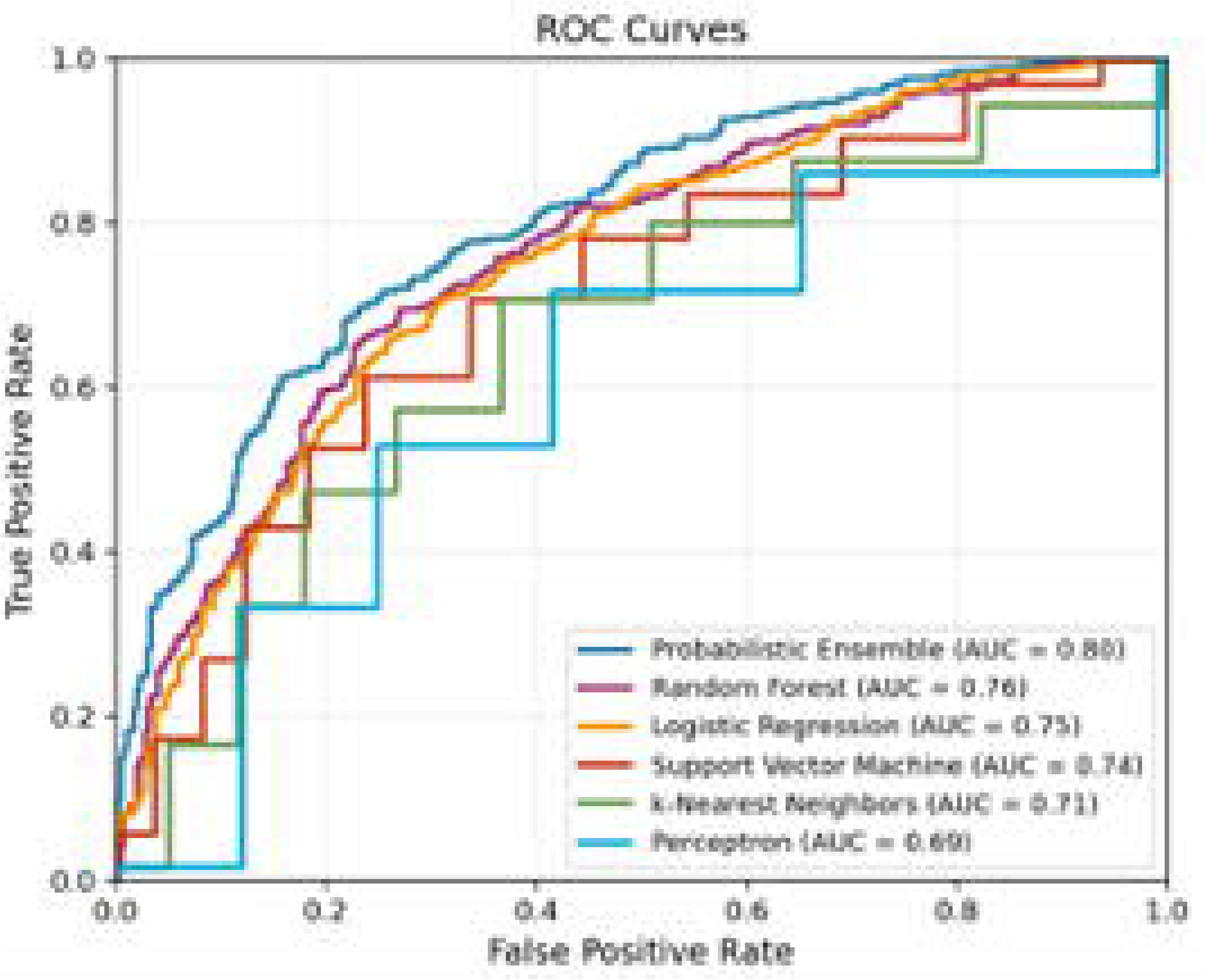
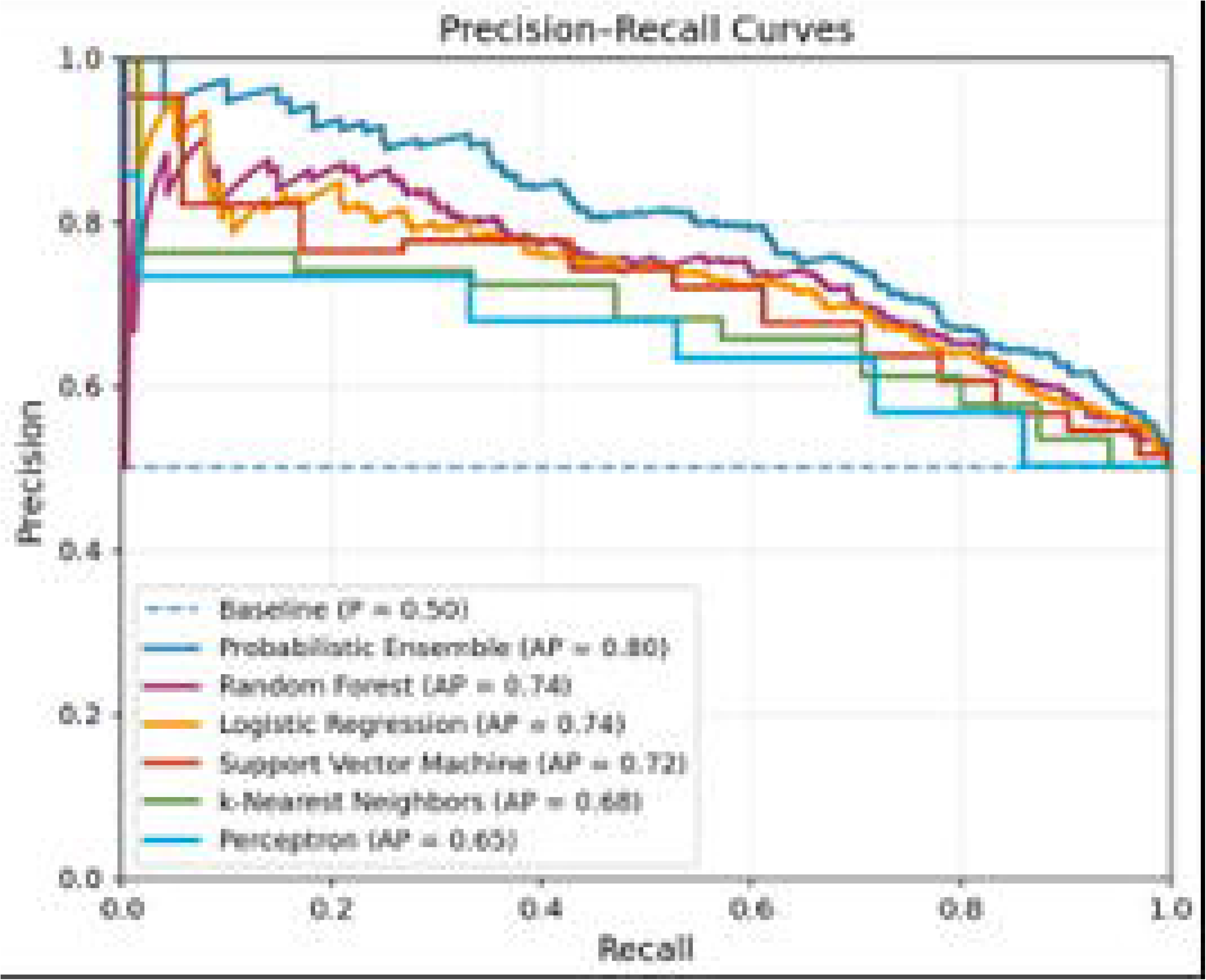
Classification performance for CE versus NCE awakenings across supervised models. (Panel A) Receiver-operating-characteristic curves and ROC–AUC for each classifier. (Panel B) Precision–recall curves and average precision (AP) for each classifier. The probabilistic ensemble achieved the highest overall discrimination.

### Stability across cross-validation folds

Model performance variability across the 5 participant-level cross validation folds is summarized in Figure 4. The probabilistic ensemble demonstrated the most consistent performance across folds for ROC-AUC, precision, and recall relative to individual learners (Figure 4). Variability was more pronounced among individual classifiers, with performance distributions spanning a broader range across folds.

**Figure 4.**
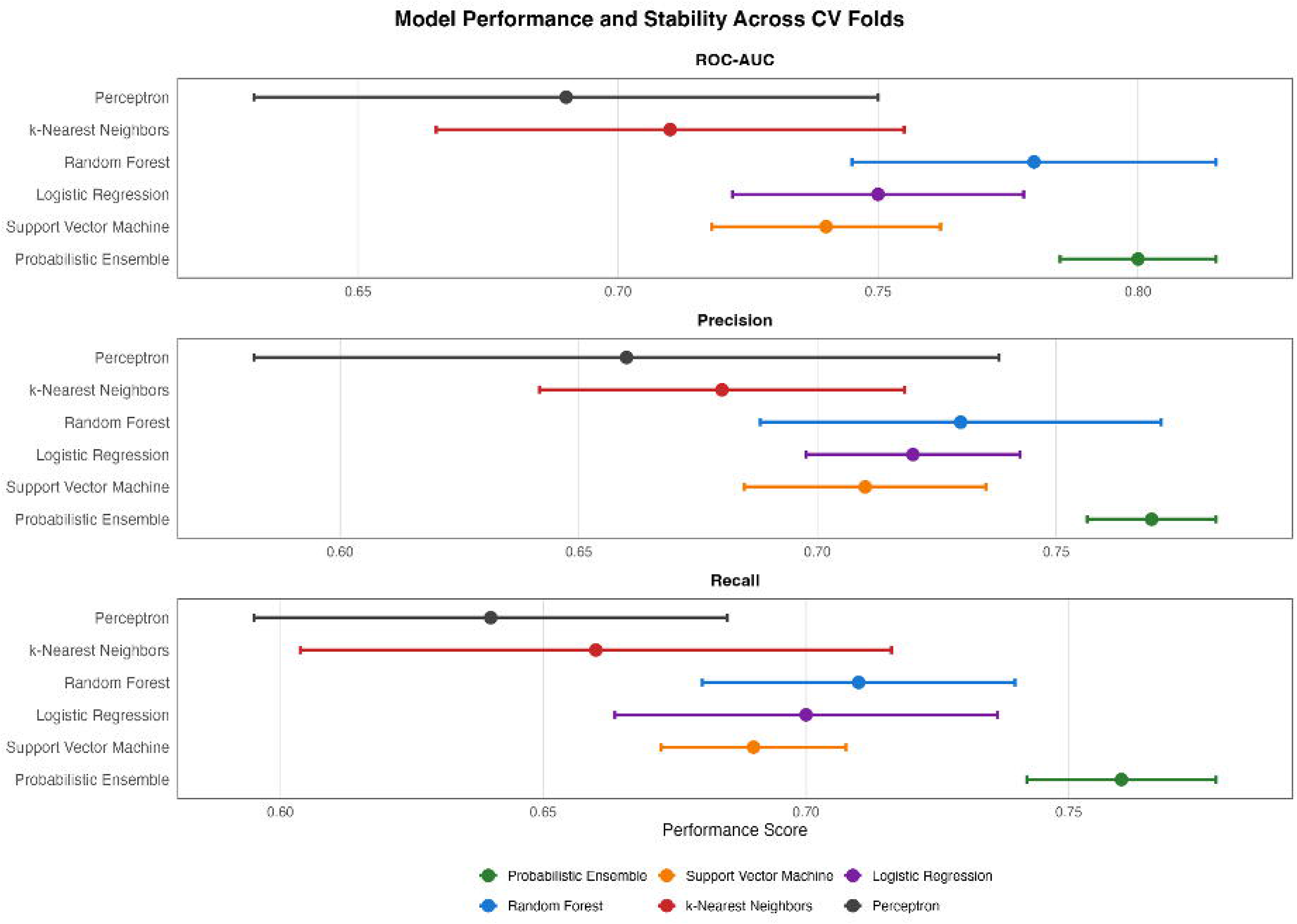
Model stability across participant-level cross-validation folds. Distributions of performance metrics across the five participant-level folds are shown for each model (ROC–AUC, precision, and recall), highlighting variability in generalization performance across splits.

## Discussion

In this serial-awakening HD-EEG study during NREM stages 2 and 3, we analyzed 699 awakenings (351 CE, 348 NCE), averaging 4.99 awakenings per participant. Across awakenings, CE and NCE differed in posterior spectral features, cross-frequency balance, and large-scale network organization. The strongest discriminators were posterior alpha and delta power, posterior gamma power, and gamma/alpha ratios, each surviving FDR correction with moderate-to-large effect sizes. Connectivity-derived measures also differed between CE and NCE, including modularity (Q), global efficiency, and rich-club coefficient, indicating that separability extended beyond regional power to network-level organization. Using participant-level cross-validation, classification performance was consistently above chance across model families, with best performance from a probabilistic ensemble (ROC-AUC 0.80, AP 0.80).

Participant-level cross-validation is particularly important in serial-awakening datasets because it reduces subject leakage and yields a more conservative estimate of generalization.(4,22) More broadly, these results reinforce that sleep consciousness can be studied in NREM using within-subject awakenings and objective electrophysiologic features rather than assuming dreaming is confined to REM sleep.(3,9,23)

### Local Posterior Dynamics and Cross-Frequency Balance

The prominence of posterior spectral differences in CE versus NCE fits well with prior work linking dream reports to changes in posterior cortical activity. Across studies, dreaming has been associated with reduced low-frequency power and increased higher-frequency activity in posterior regions thought to support perceptual content.(2,23,24) Our results are consistent with that general pattern, with strong CE–NCE separation in posterior alpha and delta alongside posterior gamma and gamma/alpha balance metrics. Because dreaming outside REM sleep is common, NREM2 and NREM3 predictors are particularly relevant for understanding how experience varies throughout the night.(3,9)

At the same time, spectral power and cross-frequency ratios can reflect several overlapping processes, including arousal fluctuations around awakenings, so these features are best framed as correlates of reportable experience rather than unique signatures of consciousness.(23,25,26) This distinction matters because CE/NCE labels depend on what participants can report after awakening, and reportability can be influenced by sleep inertia, attention to the task, and memory.(4,26) Related work using microstate and state-dynamics approaches suggests that dreaming is associated with organized spatiotemporal patterns, which complements purely spectral descriptions. Taken together, our posterior findings support a conservative interpretation: CE is associated with a physiological state characterized by shifts in posterior spectral balance and local activation, increasing the likelihood of reportable experience.(2,3,10,27)

### Network Organization: integration and segregation

In addition to regional power, CE and NCE differed in graph-theoretic measures derived from functional connectivity, indicating that experience-related differences also show up at the level of network organization.

Modularity (Q) and global efficiency are widely used summary measures that capture complementary aspects of segregation and integration in brain networks. Large-scale network analyses routinely use these measures to summarize distributed organization across brain states, supporting their use as interpretable global descriptors alongside regional features.(28,29)

Our observation that modularity was lower and global efficiency was higher during CE compared to NCE aligns with broader work showing that large-scale functional organization changes across the NREM cycle and tracks slow electrophysiologic rhythms.(34,35) At a conceptual level, this supports the idea that experience during sleep relates not only to where activity occurs but also to how distributed interactions are organized across regions.(3,32) However, network estimates can be sensitive to analytic choices, including thresholding and parameter selection, which can affect the stability of global graph metrics. For EEG-derived graphs in particular, variability introduced by thresholding and preprocessing choices is a known concern, so converging evidence across feature families strengthens interpretation.(28,33,34)

In our data, network effects appeared alongside robust posterior spectral and cross-frequency differences, supporting the view that separability is not an artifact of a single analytic family. These results also motivate stage-aware follow-up analyses, since NREM2 and NREM3 can differ in synchrony and bi-stability, which may shape both connectivity estimates and reportability.(31,35)

### Interpretation of Classification Performance

Across supervised models, discrimination ranged from ROC-AUC 0.69 (perception) to 0.80 (probabilistic ensemble), with intermediate performance for random forest (0.76), logistic regression (0.75), SVM (0.74), and KNN (0.71). Precision recall results showed the same ordering, with AP 0.80 for the ensemble and AP 0.65-0.74 across the remaining models. That linear and kernel-based methods performed competitively suggest that discriminative information is present in the selected feature space rather than depending on a single highly flexible model.

The ensemble advantage is consistent with the general ML principle that combining complementary decision rules can improve robustness in heterogeneous datasets.(22) Performance remaining below ceiling supports a conservative interpretation that scalp EEG contains probabilistic information about reported experience but does not provide a deterministic signature that generalizes perfectly across participants and awakenings.(4) This is plausible in serial-awakening paradigms, where inter-individual variability, within-subject state fluctuations, and reportability constraints can limit maximal separability. (3,26)

Because arousal level can strongly modulate electrophysiologic features near awakenings, separating experience from arousal remains an important interpretive boundary for classifiers trained on scalp EEG. (25,36) Finally, the fact that conclusions were consistent across ROC and precision–recall metrics reduces the likelihood that results depend on a single metric choice or operating threshold(22)

Several limitations should be considered when interpreting these findings. First, conscious experience was determined from brief, retrospective self-report after awakening and may underestimate the presence or richness of subjective experience, particularly when recall is incomplete. Second, the analyses were restricted to NREM stages 2 and 3, therefore these results may not generalize the awakenings from REM sleep where neurophysiology and dream phenomenology may differ, or to other states of altered consciousness such as anesthesia or after severe brain damage. Future studies may assess the generalization of our findings in such states Third, scalp HDEEG provides indirect, correlational measurements of neuronal activity. In addition, although lagged coherence was used to reduce zero-lag coupling, connectivity estimates from scalp EEG remain sensitive to volume conduction, referencing choices, and signal to noise variability. However, scalp EEG measures are more widely scalable than source-reconstructed ones. Fourth, classification performance should be interpreted as evidence of separability between labeled states rather than deterministic “signature” of consciousness, model performance may be influenced by inter-individual differences, variability in arousal state, and measurement noise. Finally, while participant-level cross-validation reduces subject leakage, external validation in independent datasets will be necessary to establish generalizability beyond the current sample and recording conditions.

Future work could extend this approach in several ways. Stage-specific analyses could test whether discriminative features and model performances differ between NREM stage 2 and 3 awakenings, and whether similar patterns generalize to REM sleep, anesthesia or unresponsive patients with severe brain damage. Incorporating additional behavioral and phenomenological measures, such as ratings, content richness, or graded reports of experience, may reduce label ambiguity and clarify whether models distinguish conscious experience itself versus recall quality. Methodologically, validating these features and models in external cohorts and across recording systems will be important for assessing robustness. Additional model analyses, including calibration assessments and threshold based operating points (sensitivity and specificity), could help translate discriminability metrics into more interpretable performance summaries. Finally, exploring reduced-channel implementations and real-time pipelines could evaluate whether comparable performance can be achieved with fewer electrodes while preserving participant-level generalization.

This study examined EEG differences between conscious experience (CE) and no conscious experience (NCE) during NREM stage 2 and 3 sleep using, spectral, connectivity and engineered cross-frequency features with supervised classification. CE and NCE were associated with differences in posterior spectral features, cross-frequency balance, and large-scale network organization, and models discriminated CE from NCE above chance, with the best performance observed for an ensemble approach. Together, these findings support the presence of reproducible, probabilistic EEG patterns associated with reported conscious experience during sleep that can predict dreaming on an individual participant level. External validation and stage-specific analyses, along with approaches that better address causality and subcortical contributions, will be important next steps.

## Methods

The institutional review board (IRB) of the University of Wisconsin (Health Sciences IRB – ID 2018-0320) approved all study procedures and all subjects provided written informed consent before participating in the study. All procedures complied with relevant guidelines and regulations. All data were de-identified prior to analysis, and no protected health information (PHI) was included in the analytic dataset.

### Participants

A total of 140 participants (male = 82 [58.57%], female = 58 [41.43%]) were included in the study. Participants ranged from 25 to 66 years of age, with a mean of 44.24 years. The racial composition of the sample was as follows: 4 Native American, 12 Asian, 2 Black, 1 Pacific Islander, and 121 White participants. Some participants identified with different racial compositions. A total of 6 identified themselves as multiple races. Participants underwent overnight high-density electroencephalography (HD-EEG) recordings using a serial awakening protocol. Throughout the night, participants were awakened by an auditory stimulus. Upon awakening, EEG data from one minute prior to awakening was extracted, and the participants were asked to report whether they had been dreaming. Only awakenings occurring from Non-Rapid Eye Movement (NREM) sleep stages 2 and 3 were included in the analyses, awakenings from NREM stage 1, NREM stage 4, and Rapid Eye Movement (REM) sleep were excluded. Awakenings were labeled as conscious experience (CE) if dream content could clearly be reported and as no conscious experience (NCE) if the participant indicated they were not dreaming and unable to recall dream content. In total, 699 awakenings met inclusion criteria and were analyzed (351 CE and 348 NCE).

### Regional Definitions

To enable an anatomically informed spectral analysis, we began by grouping electrodes into posterior, frontal, and control regions. Posterior regions (Left: [‘TP7’, ‘P7’, ‘P5’, ‘P3’, ‘P1’, ‘PPO5’, ‘PPO3’, ‘PO7’, ‘PO3’] Midline: [‘Pz’, ‘PPOz’, ‘POz’, ‘Oz’, ‘CPPz’, ‘Iz’] Right: [‘TP8’, ‘P8’, ‘P6’, ‘P4’, ‘P2’, ‘PPO6’, ‘PPO4’, ‘PO8’, ‘PO4’]) primarily include parieto-occipital sites. Frontal regions ([‘F7’, ‘F5’, ‘F3’, ‘F1’, ‘AF7’, ‘AF5’, ‘AF3’, ‘AF1’, ‘Fp1’] Midline: [‘Fz’, ‘AFz’, ‘Fpz’, ‘FCz’, ‘FFCz’, ‘FFC1h’] Right: [‘F8’, ‘F6’, ‘F4’, ‘F2’, ‘AF8’, ‘AF6’, ‘AF4’, ‘AF2’, ‘Fp2’]) included anterior and prefrontal electrodes, and control (Left: [‘C3’, ‘CP3’] Midline: [‘Cz’, ‘CPz’, ‘FCz’] Right: [‘C4’, ‘CP4’]) regions consisted of central electrodes less consistently associated with consciousness-specific findings, and aim to act as a baseline for regional comparisons (Figure 1).

### Power Spectral Analysis

Power spectral density (PSD) was computed using Welch’s method (37) with overlapping windows (2-s windows 50% overlap). Band-limited power was extracted for canonical frequency bands: delta (0.5-4 Hz), theta (4-8 Hz), alpha (8-12 Hz), beta (12-30 Hz), and gamma (30-40 Hz). (38,39) Regional band power was obtained by averaging PSD values across electrodes within each anatomical grouping.

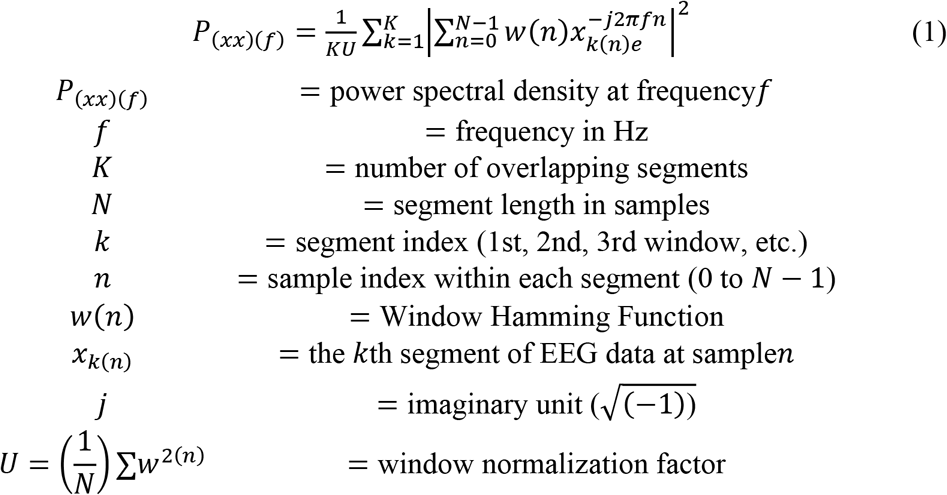

### Functional Connectivity

To quantify functional connectivity, we computed adjacency matrices using Lagged Coherence, a popular phase-based metric designed to minimize zero-lag coupling and volume conduction artifacts. Lagged coherence between two signals was defined as the squared imaginary component of the cross-spectral density normalized by the power spectra of individual signals.

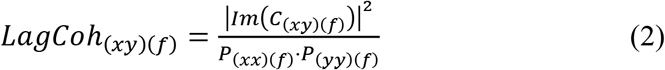

We employed a sliding-window approach (2-s windows, 50% overlap) to compute adjacency matrices for each awakening across canonical frequency bands: Delta (0.5–4 Hz), Theta (4–8 Hz), Alpha (8–12 Hz), Beta (12–30 Hz), and Gamma (30–40 Hz). Each awakening contributed to its own set of windowed adjacency matrices. Windowed matrices were averaged within band to produce one connectivity matrix per band per awakening.

### Graph-Theoretic Network Measures

Weighted adjacency matrices were then used to derive global graph-theoretic (GT) metrics characterizing network organization. Three global measures arose as discriminant. Global efficiency, indexing network integration by using the inverse shortest path lengths between nodes. Modularity (Q), indexing community structures, and network segregation. Rich-club coefficient, indexing preferential connectivity among high-degree nodes.

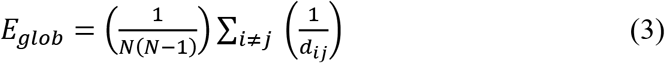

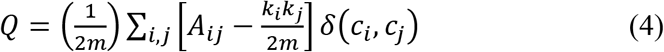

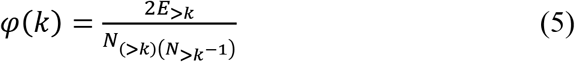

These measures capture the complementary aspects of network organization and are conceptually consistent with theoretical frameworks emphasizing large-scale information integration in conscious states

### Engineered Features

In addition to spectral and connectivity features, additional features were created to enhance discriminative power while ensuring no extraneous features were introduced to the model. Two features arose as highly significant, and uncorrelated with the existing feature set. These include gamma/alpha power ratios in both the posterior and frontal region. These features are engineered to capture cross-frequency associated with conscious processing while ensuring the absence of multicollinearity.

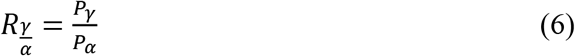

### Feature Pool and Statistical Screening

Approximately 60 candidate features spanning spectral, connectivity and engineered domain were initially computed. Group differences between CE and NCE awakenings were assessed using Mann-Whitney U tests. P-values were then corrected using the Benjamini-Hochberg false discovery rate (FDR) procedure. (40) In order to better emphasize meaningful differences, features were only retained if and only if they demonstrated moderate or large effect sizes evidenced by a |Rank-Biserial| > 0.4. This step helped us to find group separation beyond statistical fluctuations.

### Nonlinear Dependency and Multicollinearity Control

Neural features often exhibit nonlinear relationships and inter-feature correlations; mutual information analysis was used to quantify feature-label dependencies without assuming linearity. LASSO regularization was subsequently applied to reduce redundancy and mitigate multicollinearity, creating a parsimonious subset of strong, predictive features.

### Machine Learning Models

The selected features were then used as inputs to several supervised learning algorithms, including logistic regression, support vector machines (SVM), feed-forward neural networks, k-nearest neighbors, linear classifiers, and ensemble methods including random forests, and a probabilistic ensemble. This diversity allows us to compare the effectiveness between linear, nonlinear, kernel, and ensemble-based approaches. For each model, model performance was evaluated using 5-fold cross-validation grouped at the participant level. All awakenings from a given participant were assigned to the same fold to prevent subject leakage and over-representation. Folds were constructed to balance the participants and maintain class distribution whenever possible. Performance metrics were computed on held-out folds and averaged across splits to estimate the respective model’s generalization performance. Primary evaluation focused on ROC-AUC due to its robustness to classification thresholds and class imbalances.

## Supporting information

Supplemental Figure 1, Experimental Design

## Glossary

HDEEG: High Density Electroencephalography
NREM: Non-Rapid Eye Movement
REM: Rapid Eye Movement
NCE: Non-Conscious Experience
CE: Conscious Experience
ROC-AUC: Receiver Operating Curve-Area Under Curve
IRB: Institutional Review Board
PHI: Protected Health Information
PSD: Power Spectral Density
GT: Graph Theoretic
SVM: Support Vector Machine
FDR: False Discovery Rate
KNN: K-Nearest Neighbor

## Notes

### Competing Interest Statement

The authors have declared no competing interest.

## References

1. Tononi G, Boly M, Cirelli C. Consciousness and sleep. Neuron. 2024 May;112(10):1568–94. doi:10.1016/j.neuron.2024.04.011

2. Siclari F, Baird B, Perogamvros L, Bernardi G, LaRocque JJ, Riedner B, et al. The neural correlates of dreaming. Nat Neurosci. 2017 Jun;20(6):872–8. doi:10.1038/nn.4545

3. Nir Y, Tononi G. Dreaming and the brain: from phenomenology to neurophysiology. Trends Cogn Sci. 2010 Feb;14(2):88–100. doi:10.1016/j.tics.2009.12.001

4. Siclari F, LaRocque JJ, Postle BR, Tononi G. Assessing sleep consciousness within subjects using a serial awakening paradigm. Front Psychol. 2013;4. doi:10.3389/fpsyg.2013.00542

5. Kahn D, Gover T. Consciousness In Dreams. In: International Review of Neurobiology [Internet]. Elsevier; 2010 [cited 2026 Mar 4]. p. 181–95. Available from: https://linkinghub.elsevier.com/retrieve/pii/S0074774210920096 doi:10.1016/S0074-7742(10)92009-6

6. Diezig S, Denzer S, Achermann P, Mast FW, Koenig T. EEG Microstate Dynamics Associated with Dream-Like Experiences During the Transition to Sleep. Brain Topogr. 2024 Mar;37(2):343–55. doi:10.1007/s10548-022-00923-y

7. Siclari F, Bernardi G, Cataldi J, Tononi G. Dreaming in NREM Sleep: A High-Density EEG Study of Slow Waves and Spindles. J Neurosci. 2018 Oct 24;38(43):9175–85. doi:10.1523/JNEUROSCI.0855-18.2018

8. Prerau MJ, Brown RE, Bianchi MT, Ellenbogen JM, Purdon PL. Sleep Neurophysiological Dynamics Through the Lens of Multitaper Spectral Analysis. Physiology. 2017 Jan;32(1):60–92. doi:10.1152/physiol.00062.2015

9. Zhang J, Wamsley EJ. EEG predictors of dreaming outside of REM sleep. Psychophysiology. 2019 Jul;56(7):e13368. doi:10.1111/psyp.13368

10. Bréchet L, Brunet D, Perogamvros L, Tononi G, Michel CM. EEG microstates of dreams. Sci Rep. 2020 Oct 13;10(1):17069. doi:10.1038/s41598-020-74075-z

11. Schiff ND. Posterior medial corticothalamic connectivity and consciousness. Ann Neurol. 2012 Sep;72(3):305–6. doi:10.1002/ana.23671

12. Koessler L, Maillard L, Benhadid A, Vignal JP, Felblinger J, Vespignani H, et al. Automated cortical projection of EEG sensors: Anatomical correlation via the international 10–10 system. NeuroImage. 2009 May 15;46(1):64–72. doi:10.1016/j.neuroimage.2009.02.006

13. Schartner M, Seth A, Noirhomme Q, Boly M, Bruno MA, Laureys S, et al. Complexity of Multi-Dimensional Spontaneous EEG Decreases during Propofol Induced General Anaesthesia. Chialvo DR, editor. PLOS ONE. 2015 Aug 7;10(8):e0133532. doi:10.1371/journal.pone.0133532

14. Ma Y, Hamilton C, Zhang N. Dynamic Connectivity Patterns in Conscious and Unconscious Brain. Brain Connect. 2017 Feb;7(1):1–12. doi:10.1089/brain.2016.0464

15. Luppi AI, Craig MM, Pappas I, Finoia P, Williams GB, Allanson J, et al. Consciousness-specific dynamic interactions of brain integration and functional diversity. Nat Commun. 2019 Oct 10;10(1):4616. doi:10.1038/s41467-019-12658-9

16. Crone JS, Schurz M, Höller Y, Bergmann J, Monti M, Schmid E, et al. Impaired consciousness is linked to changes in effective connectivity of the posterior cingulate cortex within the default mode network. NeuroImage. 2015 Apr;110:101–9. doi:10.1016/j.neuroimage.2015.01.037

17. Koch C, Massimini M, Boly M, Tononi G. Neural correlates of consciousness: progress and problems. Nat Rev Neurosci. 2016 May;17(5):307–21. doi:10.1038/nrn.2016.22

18. Fell J, Axmacher N, Haupt S. From alpha to gamma: Electrophysiological correlates of meditation-related states of consciousness. Med Hypotheses. 2010 Aug;75(2):218–24. doi:10.1016/j.mehy.2010.02.025

19. Albantakis L, Barbosa L, Findlay G, Grasso M, Haun AM, Marshall W, et al. Integrated information theory (IIT) 4.0: Formulating the properties of phenomenal existence in physical terms. Graham LJ, editor. PLOS Comput Biol. 2023 Oct 17;19(10):e1011465. doi:10.1371/journal.pcbi.1011465

20. Baars BJ. Global workspace theory of consciousness: toward a cognitive neuroscience of human experience. In: Progress in Brain Research [Internet]. Elsevier; 2005 [cited 2026 Mar 4]. p. 45–53. Available from: https://linkinghub.elsevier.com/retrieve/pii/S0079612305500049 doi:10.1016/S0079-6123(05)50004-9

21. Pascucci D, Menétrey MQ, Passarotto E, Luo J, Paramento M, Rubega M. EEG brain waves and alpha rhythms: Past, current and future direction. Neurosci Biobehav Rev. 2025 Sep;176:106288. doi:10.1016/j.neubiorev.2025.106288

22. Moctezuma LA, Molinas M, Abe T. Unlocking Dreams and Dreamless Sleep: Machine Learning Classification With Optimal EEG Channels. Ardila CM, editor. BioMed Res Int. 2025 Jan;2025(1):3585125. doi:10.1155/bmri/3585125

23. Desseilles M, Dang-Vu TT, Sterpenich V, Schwartz S. Cognitive and emotional processes during dreaming: A neuroimaging view. Conscious Cogn. 2011 Dec;20(4):998–1008. doi:10.1016/j.concog.2010.10.005

24. Siclari F, Bernardi G, Cataldi J, Tononi G. Dreaming in NREM Sleep: A High-Density EEG Study of Slow Waves and Spindles. J Neurosci. 2018 Oct 24;38(43):9175–85. doi:10.1523/JNEUROSCI.0855-18.2018

25. Lendner JD, Helfrich RF, Mander BA, Romundstad L, Lin JJ, Walker MP, et al. An electrophysiological marker of arousal level in humans. eLife. 2020 Jul 28;9:e55092. doi:10.7554/eLife.55092

26. Marzano C, Ferrara M, Mauro F, Moroni F, Gorgoni M, Tempesta D, et al. Recalling and Forgetting Dreams: Theta and Alpha Oscillations during Sleep Predict Subsequent Dream Recall. J Neurosci. 2011 May 4;31(18):6674–83. doi:10.1523/JNEUROSCI.0412-11.2011

27. Schartner MM, Pigorini A, Gibbs SA, Arnulfo G, Sarasso S, Barnett L, et al. Global and local complexity of intracranial EEG decreases during NREM sleep. Neurosci Conscious. 2017 Jan 27;niw022. doi:10.1093/nc/niw022

28. Sporns O. Graph theory methods: applications in brain networks. Dialogues Clin Neurosci. 2018 Jun 30;20(2):111–21. doi:10.31887/DCNS.2018.20.2/osporns

29. Taberna GA, Samogin J, Zhao M, Marino M, Guarnieri R, Cuartas Morales E, et al. Large-scale analysis of neural activity and connectivity from high-density electroencephalographic data. Comput Biol Med. 2024 Aug;178:108704. doi:10.1016/j.compbiomed.2024.108704

30. Tagliazucchi E, Von Wegner F, Morzelewski A, Brodbeck V, Borisov S, Jahnke K, et al. Large-scale brain functional modularity is reflected in slow electroencephalographic rhythms across the human non-rapid eye movement sleep cycle. NeuroImage. 2013 Apr;70:327–39. doi:10.1016/j.neuroimage.2012.12.073

31. Nobili L. Local aspects of sleep: Observations from intracerebral recordings in humans. Int J Psychophysiol. 2012 Sep;85(3):356–7. doi:10.1016/j.ijpsycho.2012.06.177

32. Sarasso S, Casali AG, Casarotto S, Rosanova M, Sinigaglia C, Massimini M. Consciousness and complexity: a consilience of evidence. Neurosci Conscious. 2021 Aug 17;2021(2):niab023. doi:10.1093/nc/niab023

33. Garrison KA, Scheinost D, Finn ES, Shen X, Constable RT. The (in)stability of functional brain network measures across thresholds. NeuroImage. 2015 Sep;118:651–61. doi:10.1016/j.neuroimage.2015.05.046

34. Adamovich T, Zakharov I, Tabueva A, Malykh S. The thresholding problem and variability in the EEG graph network parameters. Sci Rep. 2022 Nov 4;12(1):18659. doi:10.1038/s41598-022-22079-2

35. Moffet EW, Verhagen R, Jones B, Findlay G, Juan E, Bugnon T, et al. Local Sleep Slow-Wave Activity Colocalizes With the Ictal Symptomatogenic Zone in a Patient With Reflex Epilepsy: A High-Density EEG Study. Front Syst Neurosci. 2020 Oct 21;14:549309. doi:10.3389/fnsys.2020.549309

36. Colombo MA, Napolitani M, Boly M, Gosseries O, Casarotto S, Rosanova M, et al. The spectral exponent of the resting EEG indexes the presence of consciousness during unresponsiveness induced by propofol, xenon, and ketamine. NeuroImage. 2019 Apr;189:631–44. doi:10.1016/j.neuroimage.2019.01.024

37. Welch P. The use of fast Fourier transform for the estimation of power spectra: A method based on time averaging over short, modified periodograms. IEEE Trans Audio Electroacoustics. 1967 Jun;15(2):70–3. doi:10.1109/TAU.1967.1161901

38. Klimesch W. The frequency architecture of brain and brain body oscillations: an analysis. Eur J Neurosci. 2018 Oct;48(7):2431–53. doi:10.1111/ejn.14192

39. Abhang PA, Gawali BW, Mehrotra SC. Technological Basics of EEG Recording and Operation of Apparatus. In: Introduction to EEG- and Speech-Based Emotion Recognition [Internet]. Elsevier; 2016 [cited 2026 Mar 4]. p. 19–50. Available from: https://linkinghub.elsevier.com/retrieve/pii/B9780128044902000026 doi:10.1016/B978-0-12-804490-2.00002-6

40. Benjamini Y, Hochberg Y. Controlling the False Discovery Rate: A Practical and Powerful Approach to Multiple Testing. J R Stat Soc Ser B Stat Methodol. 1995 Jan 1;57(1):289–300. doi:10.1111/j.2517-6161.1995.tb02031.x

