## Supplementary figures and images for "Neural Signatures of Conscious Experience During Sleep: A Serial Awakening Study Using High-Density EEG"

### Supplemental Figure 1, Experimental Design

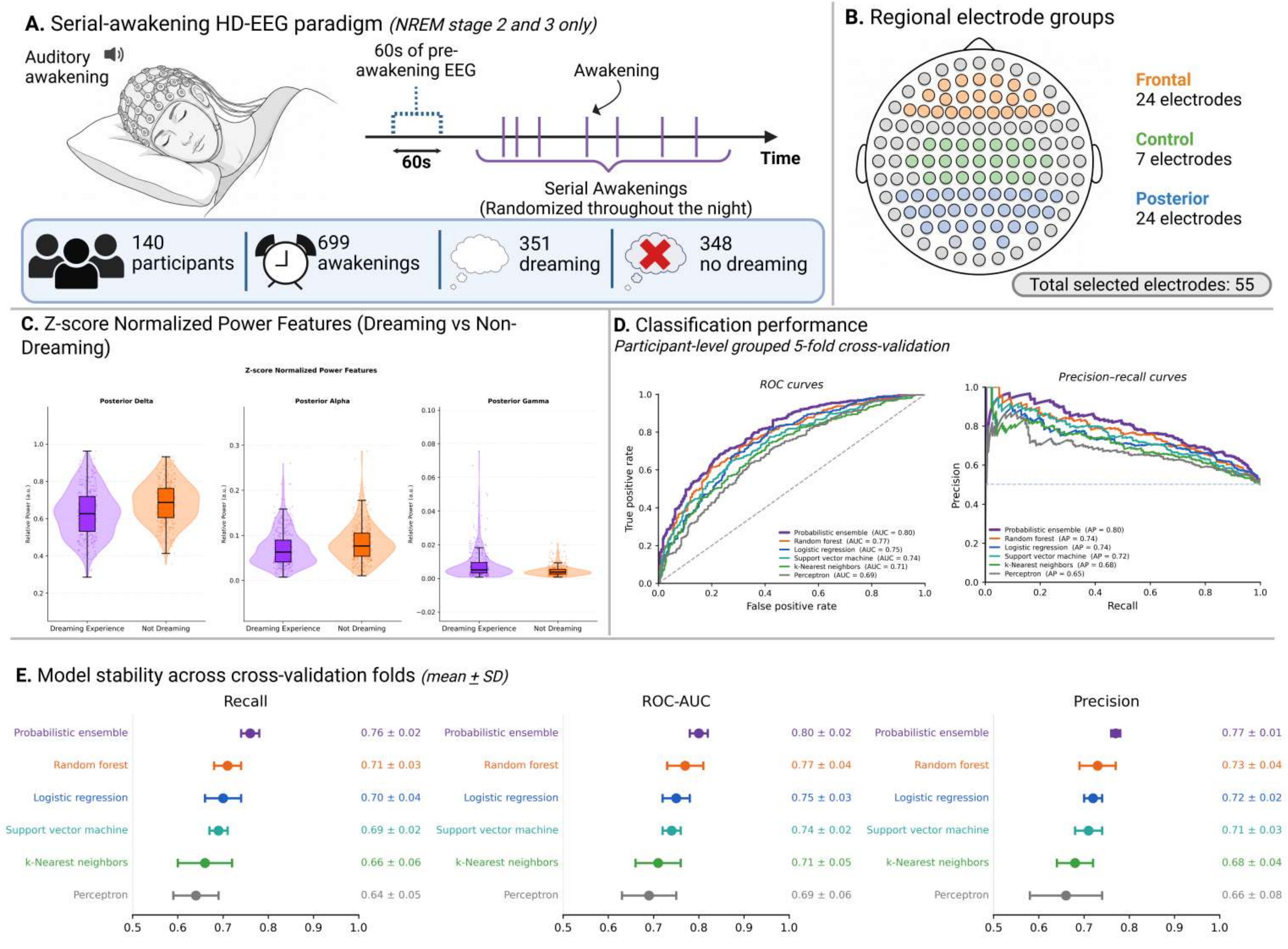


Created in https://BioRender.com
